# The kinase inhibitor AT9283 selectively kills colorectal cancer cells with hyperactive NRF2

**DOI:** 10.1101/812909

**Authors:** Laura Torrente, Gunjit Maan, Laura Casares, Angus Jackson, Tadashi Honda, Geoff Wells, Evgeny Kulesskiy, Jani Saarela, Albena T. Dinkova-Kostova, Laureano de la Vega

## Abstract

Aberrant hyperactivation of NRF2 is a common event in many tumour types and associates with resistance to therapy and poor patient prognosis. The identification of ways to overcome the protection provided by NRF2 and selectively kill cancer cells addicted to NRF2 is a desirable goal. Exploiting the CRISPR/Cas9 technology, we generated colorectal cancer cell lines with hyperactive NRF2, and used them to perform a drug screen. We identified AT9283, an Aurora kinase inhibitor, for its selectivity towards killing cancer cells with hyperactive NRF2 as a consequence to either genetic or pharmacological activation. Our results show that hyperactivation of NRF2 presents a potential vulnerability that could be therapeutically exploited, and further suggest that AT9283, a drug that is currently in clinical trials, holds promise for the treatment of tumours with hyperactive NRF2.

**Highlights:**

- We present a new model for NRF2 hyperactivation in colorectal cancer cells.
- AT9283 selectively kills cancer cells with hyperactive NRF2
- Both genetic and pharmacological activation of NRF2 sensitise cells to AT9283

## INTRODUCTION

The transcription factor NRF2 (nuclear factor erythroid 2 [NF-E2] p45-related factor 2, encoded by *NFE2L2*) is the master regulator of oxidative stress responses, which allows adaptation and survival during stress conditions. In normal cells under non-stress conditions, NRF2 levels and activity are kept low by its fast proteasomal degradation, which is principally facilitated by KEAP1 (a substrate adaptor for a Cul3-based E3 ubiquitin ligase)[1,2]. Although NRF2 is a well-characterised cytoprotective factor in normal cells, its sustained activation protects tumour cells against chemotherapy and promotes metabolic switches that support cell proliferation and tumour growth[3–6]. NRF2 hyperactivation has been observed in a variety of tumour types and is linked to poor prognosis. This sustained activation of the NRF2 pathway can be due to loss-of-function mutations in *KEAP1[3,7];* gain-of-function mutations in *NFE2L2*[7,8]; oncogenic mutations that transcriptionally induce NRF2 (i.e. KRAS, BRAF, MYC) or PTEN inactivation [9,10]); as well as epigenetic modifications leading to KEAP1 transcriptional inactivation[11,12]. Altogether, this shows that aberrant activation of the NRF2 pathway is a common event in many cancer types, and thus, the identification of ways to overcome the protection provided by NRF2 and killing cancer cells with hyperactive NRF2 is a desirable goal.

Multiple efforts have been made to identify NRF2 inhibitors and several compounds have been described to inhibit NRF2[13], but no specific inhibitor for NRF2 has been identified. Furthermore, the safety of a systemic treatment with an NRF2 inhibitor for patients is uncertain due to the critical cytoprotective role of NRF2 in normal cells and its important role in the immune response[14–16]. An alternative way to overcome the NRF2-mediated aggressiveness in cancer is to identify vulnerabilities associated with NRF2 hyperactivation. Because such approach is not expected to affect normal cells, it will be more specific and consequently safer than inhibiting NRF2 globally.

Currently, there are several models to study NRF2 hyperactivation in lung cancer, and a number of associated vulnerabilities have been identified by using KEAP1-deficient or loss-of-function mutant models[17–20]. However, in addition to NRF2, KEAP1 affects other proteins[21] and there are almost no studies addressing the role of NRF2 hyperactivation in the presence of wild-type KEAP1. To fill this gap, we generated isogenic colorectal (DLD1) cell lines harbouring gain-of-function (GOF) mutations in NRF2 using the CRISPR/Cas9 system. Using this model, we screened a panel of 528 drugs for their ability to kill DLD1 cells with hyperactive NRF2 whilst sparing DLD1 cells expressing normal levels of NRF2. From this screen, we identified AT9283, an Aurora kinase inhibitor, for its selectivity towards killing cancer cells with hyperactive NRF2. Furthermore, we validated this hit, and showed that drug sensitivity to AT9283 correlates with NRF2 levels/activity, in both a genetic model of NRF2 activation and in response to NRF2 pharmacological activation. Importantly, AT9283 is or has been in several clinical trials (clinicaltrials.gov), demonstrating the translational potential of our finding.

## MATERIAL AND METHODS

### Cell culture

All DLD1 cell lines were grown in DMEM containing 10% FBS at 37°C and 5% CO_2_. The cell lines were validated by STR profiling and were routinely tested to ensure that they were mycoplasma-negative. CRISPR-edited NRF2-knockout (NRF2-KO) and NRF2-gain-of-function (NRF2-GOF) DLD1 cell lines were produced as previously described[22].

### Antibodies and reagents

Antibody recognizing NRF2 (D1Z9C) was obtained from Cell Signaling Technology (Danvers, MA, USA). Anti-Beta-Actin antibody (C4) was from Santa Cruz Biotechnology (Dallas, Texas, USA). HRP-conjugated secondary antibodies were from Life Technologies (Carlsbad, California, USA). AT9283 was obtained purchased from ApexBio Technology (Houston, TX, USA). Dimethyl sulfoxide (DMSO) was from Sigma-Aldrich (Dorset, UK). *R,S*-sulforaphane (SFN) was purchased from LKT Laboratories (St. Paul, MN, USA). (±)-TBE-31 and HB229 were synthesized as described[23,24].

### Quantitative real time PCR (rt-qPCR)

RNA was extracted using RNeasy kit (Qiagen). RNA (500 ng) was reverse-transcribed to cDNA using Omniscript RT kit from QIAGEN (Hilden, Germany) supplemented with RNase inhibitor according to the manufacturer’s instructions. The resulting cDNA was analyzed using TaqMan Universal Master Mix II (Life Technologies, Carlsbad, CA, USA). Gene expression was determined using an Applied Biosystems 7300 Real-Time PCR system by the comparative ΔΔCT method. All experiments were performed at least in triplicates and data were normalized to the housekeeping gene HPRT1. The primers used were obtained from Thermo Fisher Scientific (Waltham, MA, USA) as follows: AKR1B10 (Hs00252524_m1), NFE2l2 (Hs00975961_g1) and HPRT1 (Hs02800695_m1)

### Focused oxidative stress pathway expression analysis

The Human Oxidative Stress RT2 Profiler PCR Array (QIAGEN) profiles the expression of 84 genes related to oxidative stress. The list of genes can be obtained from the QIAGEN webpage. The analysis was performed in triplicates, including several internal controls and housekeeping genes and analysed using the QIAGEN in-house software.

### Cell lysis protocol and western blotting

Cells were washed and harvested in ice-cold phosphate-buffered saline (PBS), lysed in RIPA buffer and sonicated. Lysates were cleared by centrifugation for 15 minutes at 4°C. The supernatant was mixed with SDS sample buffer and boiled for 5 minutes. Equal amounts of protein were separated by SDS-PAGE, followed by semi-dry blotting to a polyvinylidene difluoride membrane (PVDF, Thermo Fisher Scientific). After blocking of the membrane with 5% (w/v) TBST non-fat dry milk, primary antibodies were added. Appropriate secondary antibodies coupled to horseradish peroxidase were detected by enhanced chemiluminescence using Clarity™ Western ECL Blotting Substrate (Bio-Rad, Hercules, CA, USA).

### Drug screening

Drug sensitivity and resistance testing (DSRT)[25] was performed on DLD1 NRF2 WT and GOF cell lines in collaboration with The FIMM High Throughput Biomedicine unit at the University of Helsinki. The compound library included 528 substances consisting of conventional chemotherapeutics and a broad range of targeted oncology compounds. The compounds were dissolved in dimethyl sulfoxide or water and dispensed on 384-well plates (Corning, Corning, NY, USA). Each compound was plated at 5 concentrations covering a 10000-fold concentration range. The cells were incubated with the compounds, and after 72 h cell toxicity and viability was measured with CellToxGreen and CellTiter-Glo assays, respectively (Promega, Fitchburg, WI, USA) using a Pherastar FS (BMG Labtech, Offenburg, Germany) plate reader. The data were normalized to negative control wells (dimethyl sulfoxide only) and positive control wells containing 100 μM benzethonium chloride, which effectively kills all cells using Breeze analysis pipeline (Potdar et al, manuscript submitted). To assess quantitative drug profiles for each sample, we calculated a drug sensitivity score (DSS) based on the measured dose-response curves[26]. DSS is an integrative and robust drug response metric based on the normalized area under the curve, which considers all four curve fitting parameters in the logistic model. The complete screening data is available in Supplementary Data 1-3.

### Cell viability assay

Cells were seeded at a density of 2000 cells/well in 96-well plates, using five replicates for each treatment condition. On the next day, cells were treated with either 0.1% DMSO or different concentrations of AT9283 ranging from 10 nM to 10000 nM. Treated cells were incubated for five days at 37°C. Cell viability was determined by adding 20µl of Alamar Blue dye (Thermo Fischer scientific) to 200 µl of cell growth medium (DMEM). The plates were incubated with the dye for 3-6 hours at 37°C and the fluorescence (excitation at 570 nm and emission at 585 nm) was determined using a standard plate reader.

## RESULTS AND DISCUSSION

### Validation of a new colorectal cancer NRF2-GOF model

In order to create a physiologically relevant model of hyperactive NRF2, we used the CRISPR/Cas9 system to generate isogenic colorectal DLD1 cells with hyperactive (GOF) NRF2. Instead of targeting KEAP1, which in addition to NRF2 would also affect other KEAP1-binding partners, we used a gRNA that targets the DLG motif of endogenous NRF2. This motif is one of the two KEAP1-binding motifs within NRF2, and thus its deletion disrupts the functional interaction between NRF2 and KEAP1. Cas9-mediated DNA cleavage may result in out-of-frame DNA repair, leading to NRF2-KO clones. However, most of the cell clones in which the DNA has been repaired in-frame will result in NRF2-GOF clones (as described in [22]) due to a mutation or deletion of the KEAP1 binding sequence. These gain-of-function (GOF) mutations resemble NRF2 mutations found in some tumours[5,8,27].

Once DLD1 clones with hyperactive NRF2 (GOF) were identified, they were validated by measuring the protein levels of NRF2 and their response to the NRF2 inducer sulforaphane (SFN) (Fig 1A), and the expression levels of *NQO1* and *AKR1B10*, two well-characterised NRF2 target genes (Fig 1B). As we produced isogenic NRF2-KO and NRF2-GOF cell lines using the same gRNA, the direct comparison among the WT, KO and GOF cell lines confirms that the observed phenotypes are not artefacts due to off-target effects of the gRNA. To further characterise and validate these cell lines, we performed a pathway-focused gene expression analysis for 86 antioxidant genes, comparing NRF2-KO or NRF2-GOF versus NRF2-WT DLD1 cell lines (Fig 1C). From this analysis we concluded that a) NRF2-GOF cells have indeed upregulated NRF2 pathway, as indicated by the induction of NRF2 target genes; and b) the NRF2-GOF model is more physiologically relevant (no NRF2 knockout events in tumours have been reported) and also more sensitive than the NRF2-KO model to identify NRF2 regulated genes.

**Figure 1.**
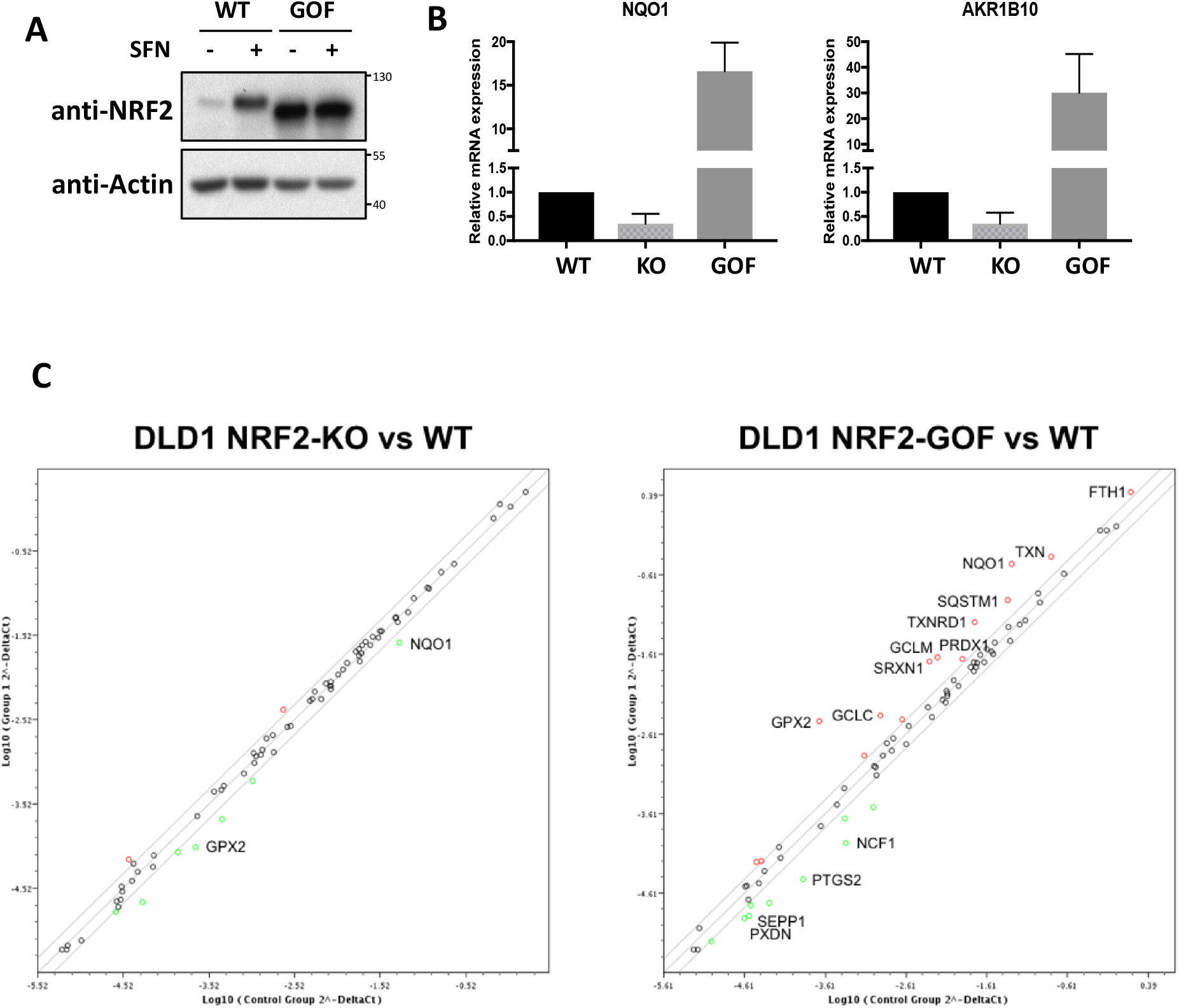
Validation of a new colorectal cancer NRF2-GOF model. **A)** Isogenic NRF2-WT and NRF2-GOF DLD1 cell lines were treated with 5 µM of SFN for 3 hours and the protein levels of NRF2 were compared. **B)** The mRNA levels for NQO1 and AKR1B10 in the different cell lines were quantified using real-time PCR. The data were normalized using β-actin as an internal control. Data represent means ± SD (n=3) and are expressed relative to the WT cells. **C)** Representation of differential expression of oxidative stress-related genes in NRF2-KO versus NRF2-WT (left panel) or NRF2-GOF versus NRF2 WT (right panel). Highlighted either in red (upregulated) or in green (downregulated) are genes with more than 2-fold change; only those with p-value< 0.05 were labelled (n=3).

### Synthetic lethality drug screening

To identify cytotoxic drugs associated to hyperactive NRF2 in colorectal cancer cell lines, we used our newly generated cell-based model. We tested 528 drugs (158 approved drugs, 285 investigational and 85 probes) for their ability to kill DLD1 cells with hyperactive NRF2 without affecting DLD1 control cells (NRF2 WT). From this screen, the drug sensitivity score (DSS), which is an integrative and robust drug response metric based on the normalized area under the curve, which considers all four curve fitting parameters in the logistic model (as explained in material and methods) was obtained (Fig 2A and Supplementary Data 1-3). Only compounds that had a difference in DSS between DLD1-GOF and DLD1-WT higher that 5 were considered positive hits. From this analysis we identified AT9283 as the top candidate with selectivity against DLD1-GOF cells (Fig 2A and 2B). AT9283 is a synthetic small heterocyclic molecule that potently inhibits several kinases, including Aurora A (3 nM), Aurora B (3 nM), JAK2 (1.2 nM), JAK3 (1.1 nM), and Abl (4.0 nM, T315I)[28]. Importantly, AT9283 has shown anti-myeloma, anti-lymphoma, anti-leukaemia and anti-colorectal cancer activity in pre-clinical studies[29–34], and its safety and efficacy against myeloma, lymphoma and leukaemia has been tested in various Phase I and II clinical trials[35–41].

**Figure 2.**
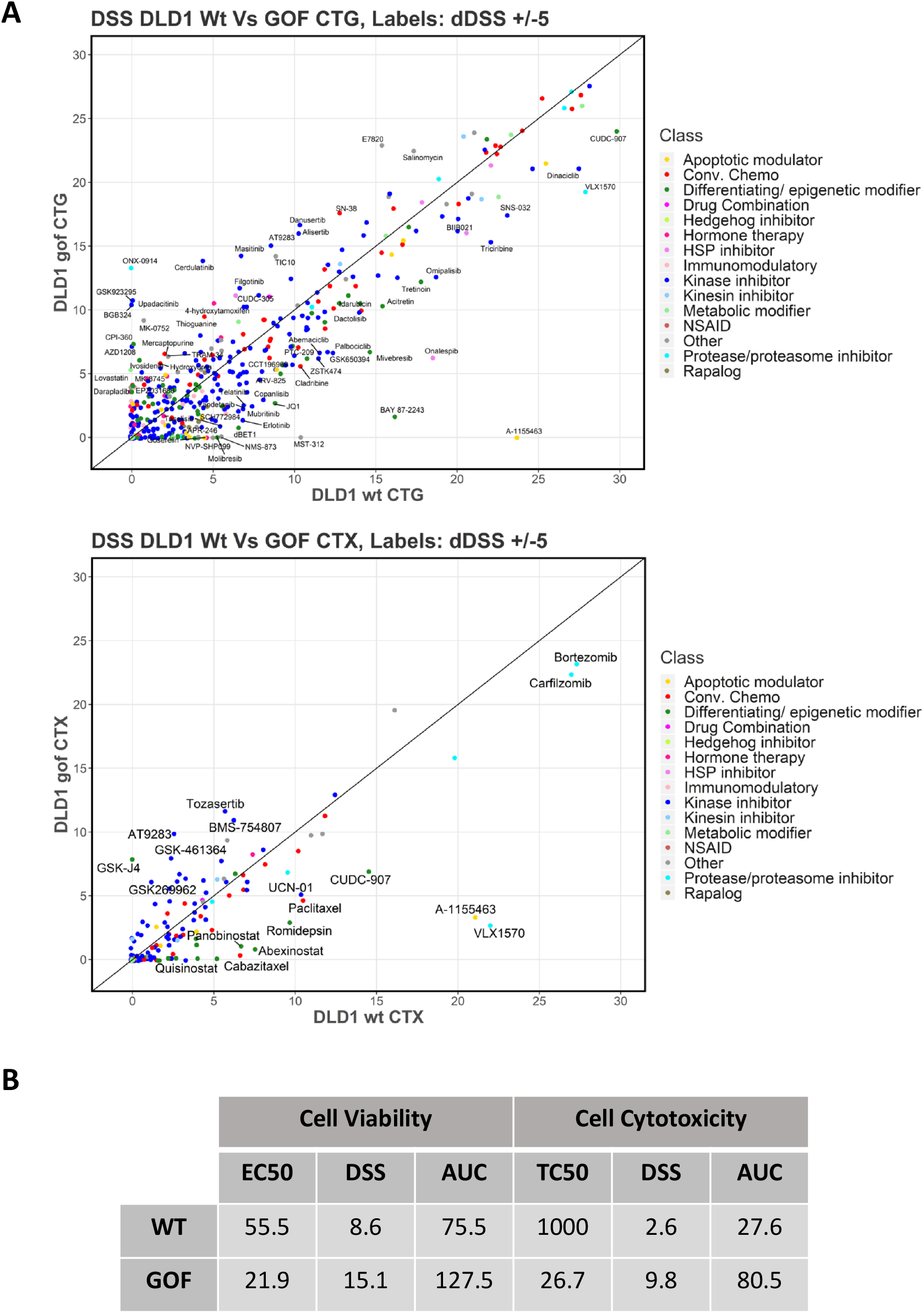
Synthetic lethality drug screening. **A)** Scatter plots showing drug sensitivity score (DSS) values of DLD1 WT and GOF cells with cell viability (CellTiter Glo, CTG) and of cell toxicity (CellTox Green, CTX) readouts. Labels are shown only for compounds which have dDSS (DSS_GOF-DSS_wt) above or below 5. **B)** Key drug response parameters of AT9283 in both WT and NRF2-GOF DLD1 cells are shown. TC_50_ (half-maximal toxic concentration), EC_50_ (half-maximal effective concentration), AUC (area under the curve), and DSS (drug sensitivity score).

### Validation of the selectivity of AT9283 against active NRF2

To validate the result from the screen, we tested the effect of AT9283 on the viability of NRF2-WT and NRF2-GOF DLD1 cells (Fig 3A). This assay confirmed the increased (by ~10-fold) sensitivity of the NRF2-GOF compared to NRF2-WT cells: AT9283 had an IC_50_ of 28 nM in NRF2-GOF cells versus an IC_50_ of 320nM in NRF2-WT cells. Further evidence that the cytotoxic effect of AT9283 was in fact dependent on the level of NRF2 activity was obtained by use of another NRF2-GOF clone with intermediate levels of NRF2 activation (GOF-Interm), as determined by NQO1 expression (Fig 3B, left panel). This analysis showed that AT9283 successfully discriminates between cell lines based on their NRF2 activity: while cells with low NRF2 activity (WT) are resistant to the compound, cells with intermediate levels of NRF2 activation (GOF Interm) are moderately sensitive to AT9283 (IC_50_ = 88 nM), and cells with high levels of NRF2 activation (GOF) are highly sensitive to AT9283. These results indicate that the activation of NRF2 in combination with nanomolar concentrations of AT9283 leads to synthetic lethality.

**Figure 3.**
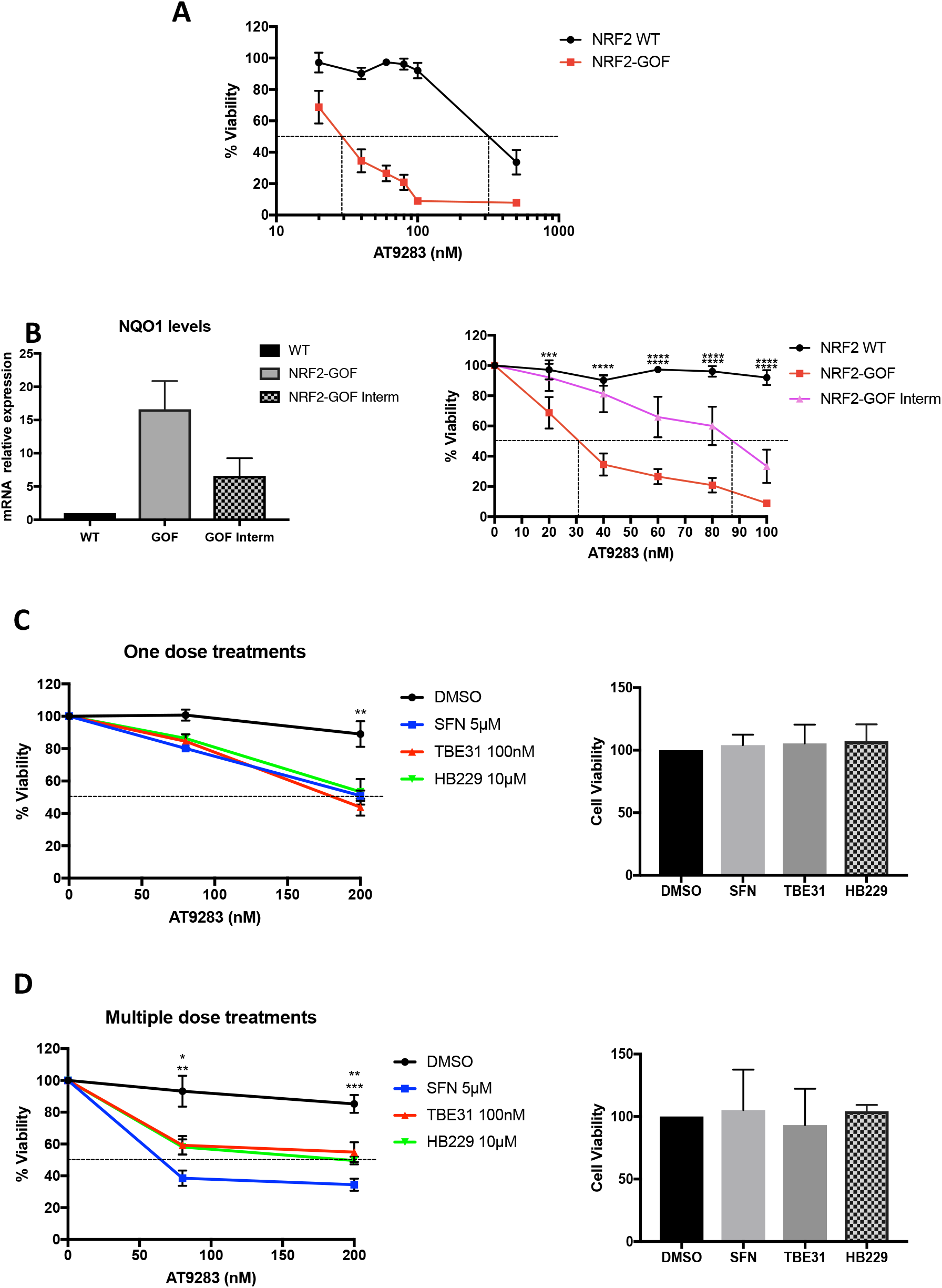
Validation of the selectivity of AT9283 against active NRF2. **A)** NRF2-WT and GOF DLD1 cells were exposed to increasing concentrations of AT9283 as indicated. After five days, cell viability was measured using Alamar Blue. Data represent means ± SD (n=4) and are expressed relative to the DMSO control which was set as 100%. **B)** The expression levels of NQO1 in NRF2-WT, NRF2-GOF and NRF2-GOF-Interm (intermediate) DLD1 cells were analysed by qRT-PCR (n=3) (left panel). NRF2-WT, NRF2-GOF and NRF2-GOF-Interm (intermediate) DLD1 cells were exposed to increasing concentrations of AT9283 as indicated. After five days, cell viability was measured using Alamar Blue (n=4). **C)** DLD1 cells were pre-treated with the indicated concentrations of NRF2 inducers or with DMSO. After 16 hours, cells were washed with PBS, and fresh media with either DMSO, 80 nM or 200 nM AT9283 was added. After three days, cell viability was measured using Alamar Blue (n=3). **D)** DLD1 cells were pre-treated with the indicated concentrations of NRF2 inducers or with DMSO as indicated. After 24 hours, fresh media with either DMSO, 80 nM or 200 nM AT9283 was added, and NRF2 inducers were replenished. On the next day, the NRF2 inducers were added again, and 24 hours later, the cell viability was measured using Alamar Blue (n=3).

To evaluate the potential therapeutic value of this synthetic lethality interaction and to further ensure that the observed effect is dependent on NRF2 activation and is not a consequence of clonal artefacts from the CRISPR-mediated GOF clones, we tested whether the selective cytotoxicity could be recapitulated by using pharmacological NRF2 activation. For this we compared the effect of three NRF2 activators that differ in potency and mechanism of action. Sulforaphane (SFN) and TBE31 are two well-characterised electrophilic compounds that react with cysteine sensors (primarily C151) of KEAP1, impairing its ability to target NRF2 for degradation[42]. By contrast, HB229 is a non-electrophilic small molecule that disrupts the KEAP1-NRF2 complex[24]. When NRF2-WT cells were pre-treated with the NRF2 activators we observed a significant sensitisation towards AT9283 (Fig 3C, left panel). Interestingly, this effect was enhanced by replenishing the NRF2 activators every 24 hours (multiple dose treatments) for the duration of the incubation period (Fig 3D, left panel). Critically, the effect on cell viability was completely dependent on AT9283 acting together with the NRF2 activator, as at these concentrations, neither AT9283 nor any of the NRF2 activators by themselves affected the viability of the cells (Fig 3C and 3D, right panels). These results demonstrate that, similar to genetic, pharmacological activation of NRF2 also sensitises DLD1 cells towards AT9283-mediated killing.

In conclusion, using a new model of a colorectal cancer cell line with hyperactive NRF2 we have identified a synthetic lethality interaction with the kinase inhibitor AT9283. Our data suggest that this drug could have clinical relevance to treat tumours with hyperactive NRF2, although further characterisation of this synthetic lethality interaction in other tumour types and in *in vivo* models is necessary to confirm its clinical potential.

## AUTHORS CONTRIBUTION

LT, GM, LC, EK and AJ conducted the experiments. LT conducted the bioinformatic analysis. LT, GM, EK and LC were responsible for initial data analysis, figure preparation and statistical analysis. AJ, TH and GW provided resources. LV and ADK had a leading contribution in the design of the study, and an active role in the discussion and interpretation of the whole dataset. LV wrote the original draft of the manuscript. ADK, LV, JS and LT reviewed and edited the manuscript. Funding acquisition LV. All the authors take full responsibility for the work.

